# cGAS, a DNA Sensor, Promotes Inflammatory Responses in Huntington Disease

**DOI:** 10.1101/2020.01.08.898718

**Authors:** Manish Sharma, Sumitha Rajendrarao, Neelam Shahani, Uri Nimrod Ramĺrez-Jarquĺn, Srinivasa Subramaniam

## Abstract

The genetic cause of Huntington disease (HD) is attributed to the N-terminal polyglutamine expansion of huntingtin (mHTT). mHTT, which is a ubiquitously expressed protein, induces noticeable damage to the striatum, which affects motor, psychiatric, and cognitive functions in HD individuals. Although inflammatory responses apparently precede striatal damage and an overall progression of HD, the molecular mechanisms at work remain unclear (1-6). In this study, we found that cyclic GMP-AMP synthase (cGAS), a DNA sensor, which regulates inflammation, autophagy, and cellular senescence (7-9), plays a critical role in the inflammatory responses of HD. Ribosome profiling analysis reveals that *cGAS mRNA* has a high ribosome occupancy at exon 1 and codon-specific pauses at positions 171 (CCG) and 172 (CGT) in HD cells, compared to the control, indicating an altered cGAS expression. Accordingly, cGAS protein levels and activity, as measured by phosphorylation of stimulator of interferon genes (STING) or TANK-binding kinase 1 (TBK1), are increased in HD striatal cells, mouse Q175HD striatum and human postmortem HD striatum, compared to the healthy control. Furthermore, cGAS-dependent inflammatory genes such as *Cxcl10* and *Ccl5* show enhanced ribosome occupancy at exon 3 and exon 1, respectively and are upregulated in HD cells. Depletion of cGAS via CRISPR/Cas-9 diminishes cGAS activity and decreases expression of inflammatory genes while suppressing the autophagy upregulation in HD cells. We additionally detected the presence of numerous micronuclei, a known inducer of cGAS, in the cytoplasm of HD cells. Overall, the data indicates that cGAS is highly upregulated in HD and mediates inflammatory and autophagy responses. Thus, targeting cGAS may offer therapeutic benefits in HD.

## Introduction

Huntington disease (HD) is a fatal neurodegenerative disorder due to huntingtin (HTT) CAG expansion mutation (coding for polyglutamine, mHTT). mHTT is a ubiquitously expressed gene, yet prominently damages the striatum and cortex, followed by widespread peripheral defects as the disease progresses. Inflammatory responses are implicated in HD. Compounds with anti-inflammatory properties have been shown to increase the survival of HD transgenic mice (10,11). Microglia, a cellular indicator of inflammation, is also increased in the striatum of HD animal and cell culture models and HD patients (12-14). Increased levels of reactive monocytes, inflammatory cytokines, chemokines, and n kynurenine/tryptophan ratio, an indicator of persistent inflammation, are all observed in pre-manifest HD patients, correlated with HD progression (15-18). Studies in mice indicate that glial cells and galectin molecules contribute to enhanced inflammatory responses in HD (19,20). Furthermore, RNA-seq analysis of tissue obtained from human HD patients and HD monkey models reveals extensive transcriptional dysregulation associated with proinflammatory pathway activation (1,21,22). Inflammation is also closely linked to autophagy, a process that is dysregulated in HD (23,24). Thus, inflammation is a prominent cellular response in HD patients and across various HD models, though the mechanisms remain less clear.

The cGAS synthase (a.k.a. Mb21d1) is an enzyme that produces cyclic guanosine monophosphate–adenosine monophosphate (cyclic GMP-AMP, cGAMP), a second messenger that activates upon binding to DNA in the cytoplasm (25). The cGAS can induce signaling that is known to promote the upregulation of inflammatory genes and play a critical role in age-related macular degeneration and cellular senescence (26-28). cGAS-induced cGAMP binds to endoplasmic reticulum-associated transmembrane protein, (STING, a. k. a TMEM173). STING recruits TANK-binding kinase 1 (TBK1), which phosphorylates transcription factors such as interferon regulatory factor 3 (IRF3), interferon regulatory factor 7 (IRF7), and other substrates such as IκB kinase α (IKKα), cRel, and p62 (sequestosome) (29,30). The cGAS synthase also plays a major role in the regulation of autophagy, suggesting a close molecular and signaling link between inflammatory response and autophagy (7,8,23,31). Despite a well-established role for cGAS in cancer, diabetes, and immune disorders, its role in neurodegenerative disease remains less clear (32-34). Upregulation of cGAMP/STING signaling, however, is linked to dopaminergic and cerebellar neuron degeneration in parkin and ataxia telangiectasia mutated ATM deficient mice, respectively (35,36). Here we investigate the role of cGAS in HD.

## Results

### Ribosome profiling data revealed high ribosome occupancy in the exon 1 on *cGAS mRNA*

For the first time, we applied high-resolution ribosome sequencing technology to genetically precise knock-in HD cell models: immortalized STHdh^Q7/Q7^ (control), STHdh^Q7/Q111^ (HD-het), and STHdh^Q111/Q111^ (HD-homo) striatal neuronal cells, derived from WT, Hdh^Q7/Q111^, and Hdh^Q111/Q111^ mouse embryos (37). We successfully generated high-quality reads and found new roles for HTT as a physiological suppressor of translational elongation by regulating ribosome movements (38). In global ribosome profiling data, we found that cGAS showed a high ribosome occupancy (ribosome protected fragments (RPF)/mRNA) estimated by ribosome protected fragments (RPF) counts in exon 1 in HD-het and HD-homo cells, while there were fewer reads in control cells (**Fig. 1A**). Ribosomes accumulated on the 5’ end of exon1 in the region (TAC CTT CTA GGC GCA TCT TCC TGC TGC) that codes for MEDPRRRTT (**inset, Fig. 1B**), indicating that *cGAS mRNA* is translationally regulated in HD. Using a pause predict software (39), we found additional single codon-pauses in 171 (CCG) (**Fig. 1C, arrow**) and 172 (CGT)) in HD cells. The RFP/mRNA analysis of other known DNA sensors such as Toll-like receptor 9, which is mostly restricted to blood and immune cells (40), is not expressed in striatal cells (38). The absence in melanoma 2 (AIM2), another cytosolic DNA sensor (41), is also induced in HD cells, but to a much smaller extent compared to cGAS (the ratio of RFP/mRNA for cGAS is 14 whereas for AIM2 it is 2.2) (**Fig. 1D**). Similarly, hnRNP-A2b1 is a newly identified sensor for viral double-stranded DNA, but not endogenous DNA, in the nucleus (41), and is expressed relatively uniformly between control and HD cells (**Fig. 1E)**. We also examined whether the ribosome profiling of two major cGAS downstream targets, STING and TBK1, are altered in HD. But we found no major differences in the distribution of ribosome occupancy or mRNA expression of STING or TBK1 between the control and HD cells (**Fig. 1F, G**). STING showed an expected pause at exon 2 (arrow), which might be involved in co-translation translocation events for proper insertion into endoplasmic reticulum (42). Together, the data suggests that *cGAS mRNA* is selectively upregulated in HD cells and shows a ribosome occupancy at exon 1, indicating translational dysregulation.

**Figure 1.**
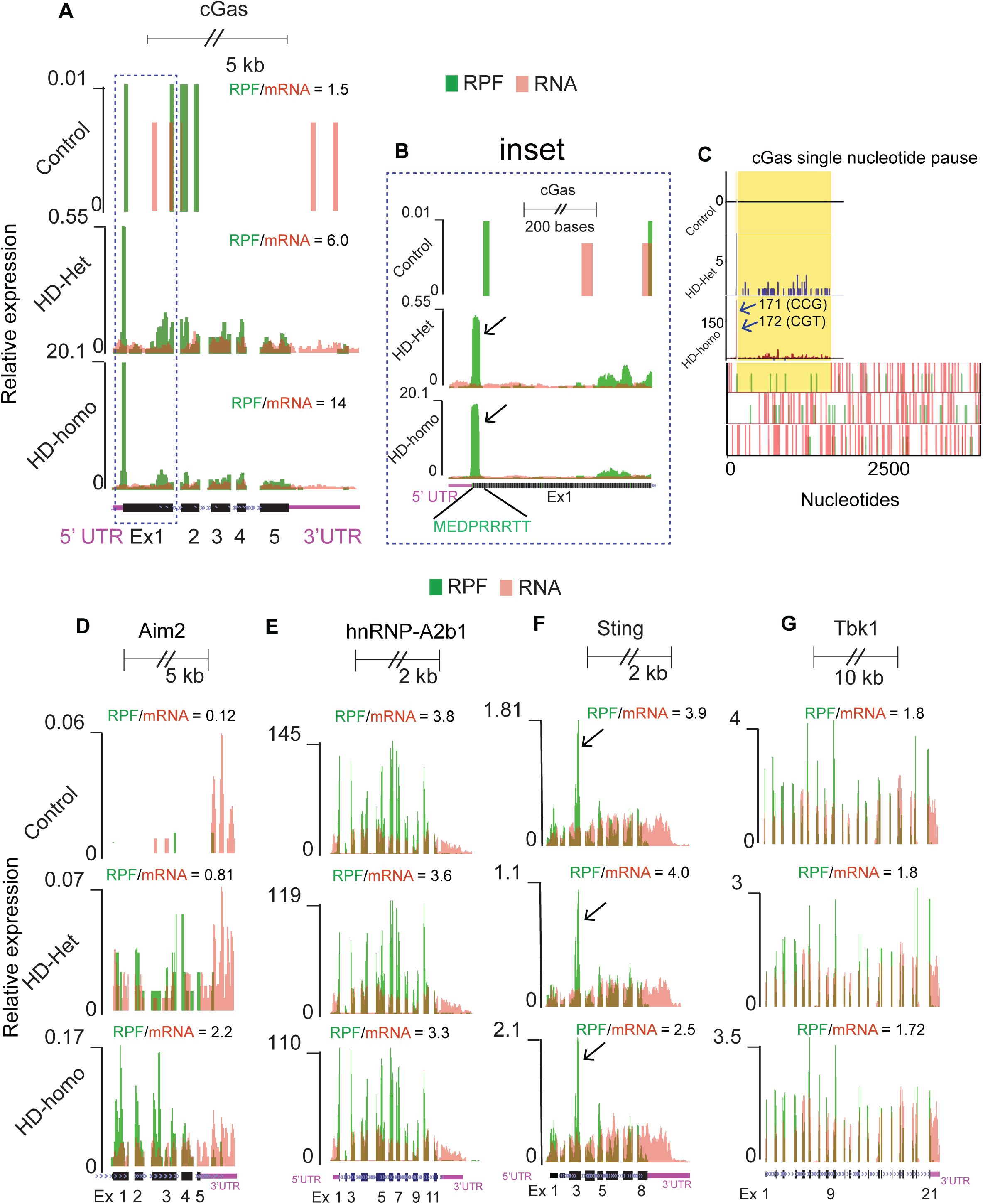
Ribosome occupancy on *cGAS* mRNA transcript in control, HD-het and HD-homo cells. (**A**) Graphs showing overlay of ribosome protected fragment (RPF)/mRNA reads for *cGAS* transcript obtained from UCSC browser. (**B**) Inset showing of exon 1 of *Htt* transcript. Arrow indicates the position of ribosome occupancy. (**C**) Graphs representing codon pause of the cGAS transcripts predicted from PausePred software from *E*. Arrow indicates the position of paused codons. (**D-G**) Graphs showing overlay of ribosome protected fragment (RPF)/mRNA reads for *hnRNP-A2b1* (D), *Aim2* (E), *Sting* (F) and *Tbk1* (G) *mRNA* transcripts obtained from UCSC browser. Arrow indicates the expected pause at Exon 2 of *Sting* mRNA due to signal peptide insertion into the endoplasmic reticulum. Ribosome footprints are shown from pooling all three replicates for control, HD-het and HD-homo cells.

### cGAS levels and its inflammatory target genes are upregulated in HD

The high ribosome occupancy of *cGAS* mRNA (**Fig. 1)** indicates that ribosomes are stalled on exon1, which may result in either increased or decreased protein production. This prompted us to investigate the protein levels of cGAS in HD cells. We found by Western blot analysis that cGAS protein is robustly upregulated in HD-het and HD-homo striatal cells, yet barely found in control striatal cells (**Fig. 2A**). This upregulation is also found in human postmortem HD patient striatum, and Q175HD-Het (neo-) striatum (43) (**Fig. 2B-C**). The data indicated that cGAS protein is upregulated in HD, and encouraged us to investigate the levels of some its known targets that are implicated in the inflammatory process (44) using our ribosome profiling data. We looked at the ribosome profiles of *Irf3, Irf7, Ccl5*, and *Cxcl10* mRNA in the HD and control cells (**Fig. 2D-G**). We found no major differences in the ribosome occupancy of *Irf3*, but a slight increase in the relative expression of *Irf7* (0.77 vs. 0.07 HD-homo vs. control), the two previously known targets of cGAS (45,46) (**Fig. 2D and E**). However, we found that *Cxcl10* and *Ccl5* mRNA are highly upregulated (30 vs 0.14 and 20 vs. 0.01 between HD Homo vs. control) in HD cells and show a high ribosome occupancy on exon 3 and exon 1 respectively, indicating translational upregulation (**Fig. 2F and G, arrow)**. Using a qPCR analysis, we further confirmed that *Ccl5* or *Cxcl10* mRNA are upregulated in HD striatal cells, HD mice striatum, as well as human HD striatum, compared to controls (**Fig. 2H I, J)**. Collectively, these data indicate that cGAS levels and its inflammatory target genes are upregulated and that their mRNAs exhibit differential ribosome occupancy in HD.

**Figure 2.**
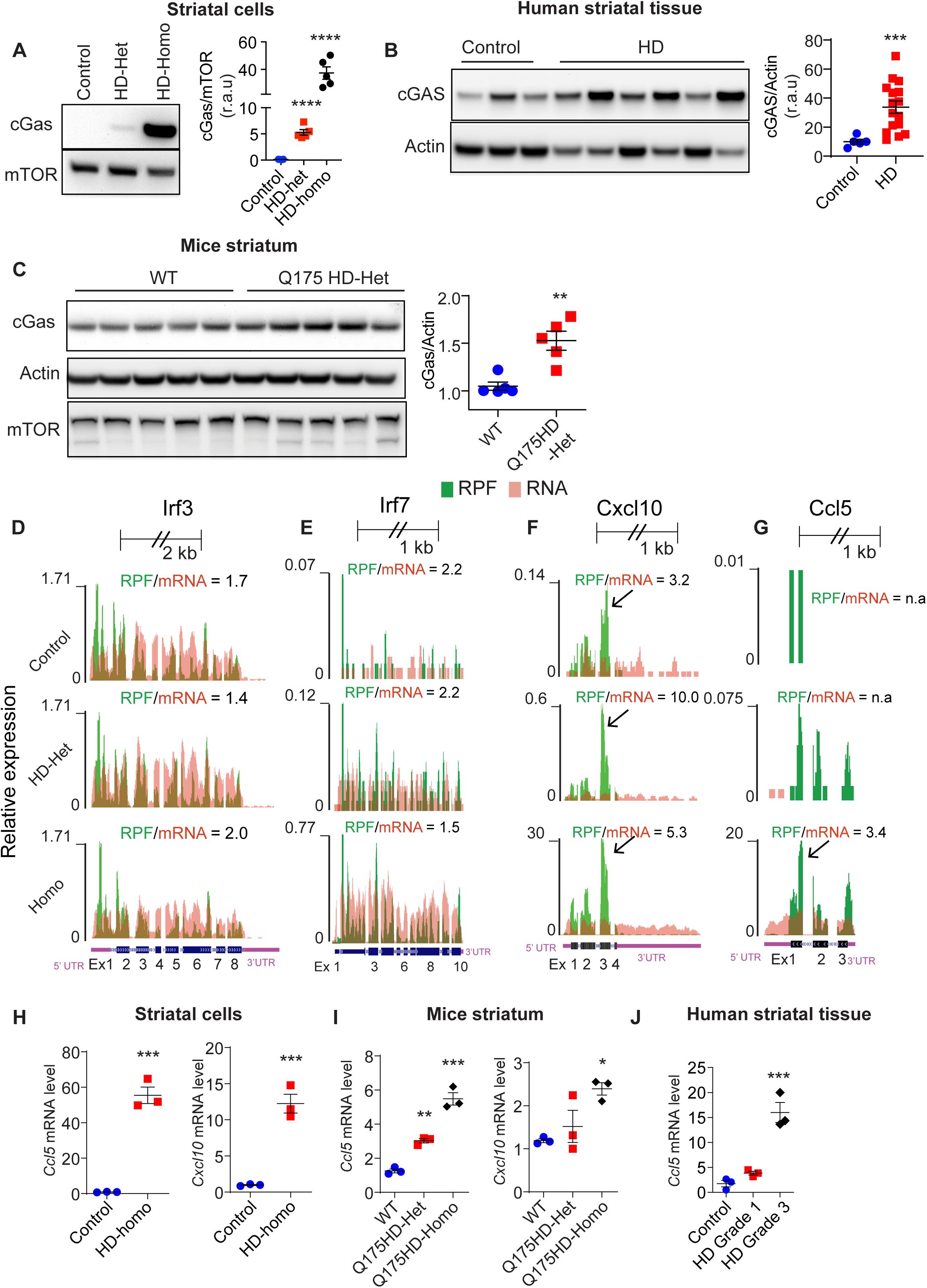
cGAS and its targets genes are upregulated in HD. (**A-D**). Representative Western blot for indicated proteins in the striatal neuronal cells (A), human unaffected (control, n = 5) and HD affected (n = 16) striatum (B), WT and Q175 HD-het (neo-) striatum (n = 5 mice/ group) (C). Error bar represents mean ± SEM, \*\**p*< 0.01, \*\*\**p*< 0.001 \*\*\*\**p*< 0.0001 by Student’s-*t* test. (**D-G**) Graphs showing overlay of ribosome protected fragment (RPF)/mRNA reads for *Irf3* (E), *Irf7* (F), *Cxcl10* (G) and *Ccl5* (H) *mRNA* transcripts obtained from UCSC browser. (**H-J**) qPCR analysis of indicated mRNA in cultured striatal cells (I), \*\*\**p*< 0.001 by Student’s-*t* test, HD mouse striatum (J) and human striatum (K). Error bar represents mean ± SEM, (n = 3), \**p*< 0.05, \*\**p*< 0.01, \*\*\**p*< 0.001 by One-Way ANOVA followed by Tukeys multiple comparison test. Ribosome footprints are shown from pooling all three replicates for control, HD-het and HD-homo cells.

### Depletion of cGAS diminishes STING and TBK1 phosphorylation and inflammatory response in HD cells

We next investigated whether cGAS upregulation is linked to the phosphorylation of its targets, the STING and TBK1 phosphorylation in control and HD cells, using Western blot analysis. We found that the phosphorylation of STING (Ser365) and TBK1 (Ser172) are increased in HD striatal cells, striatum from HD patients, and Q175HD-Het mice, compared to the control (**Fig. 3A-C**). Consistent with ribosome profiling data, we found no significant difference in the levels of IRF3, a cGAS target, or its phosphorylation (Ser-396) by Western blot analysis in striatal cells (**Fig. 3A**). We examined whether these phosphorylation events are directly controlled by cGAS. To do this, we depleted cGAS using CRISPS/Cas9 tools in control and HD-homo striatal cells. We found that upon cGAS depletion, the phosphorylation of STING and TBK1 are dramatically diminished in HD-homo cells (**Fig. 4A, B**). Note, cGAS protein is selectively diminished by ∼90% of the control in cGAS CRISPR/Cas-9, whereas mTOR, which was used an internal control, is not altered (**Fig, 4A, B**). Recent studies indicate that the cGAS/STING pathway also regulates autophagy (7,8,31), which is a major catabolic process affected in HD (24,47). We therefore investigated whether cGAS depletion also affects the autophagy HD cells. We found that autophagy, as measured by LC3B-II, is basally upregulated in HD-homo cells, compared to control cells. In cGAS depleted HD-homo cells, however, levels of LC3B-II were significantly decreased compared to control HD-homo cells (**Fig. 4C**). We further found that cGAS depleted HD-homo cells demonstrate a dramatic reduction of inflammatory gene *Cxcl10* and *Ccl5* expression (**Fig. 4D**). Collectively, this data indicates that cGAS promotes the activation of the STING/TBK1 pathway, autophagy, and inflammatory response in HD.

**Figure 3.**
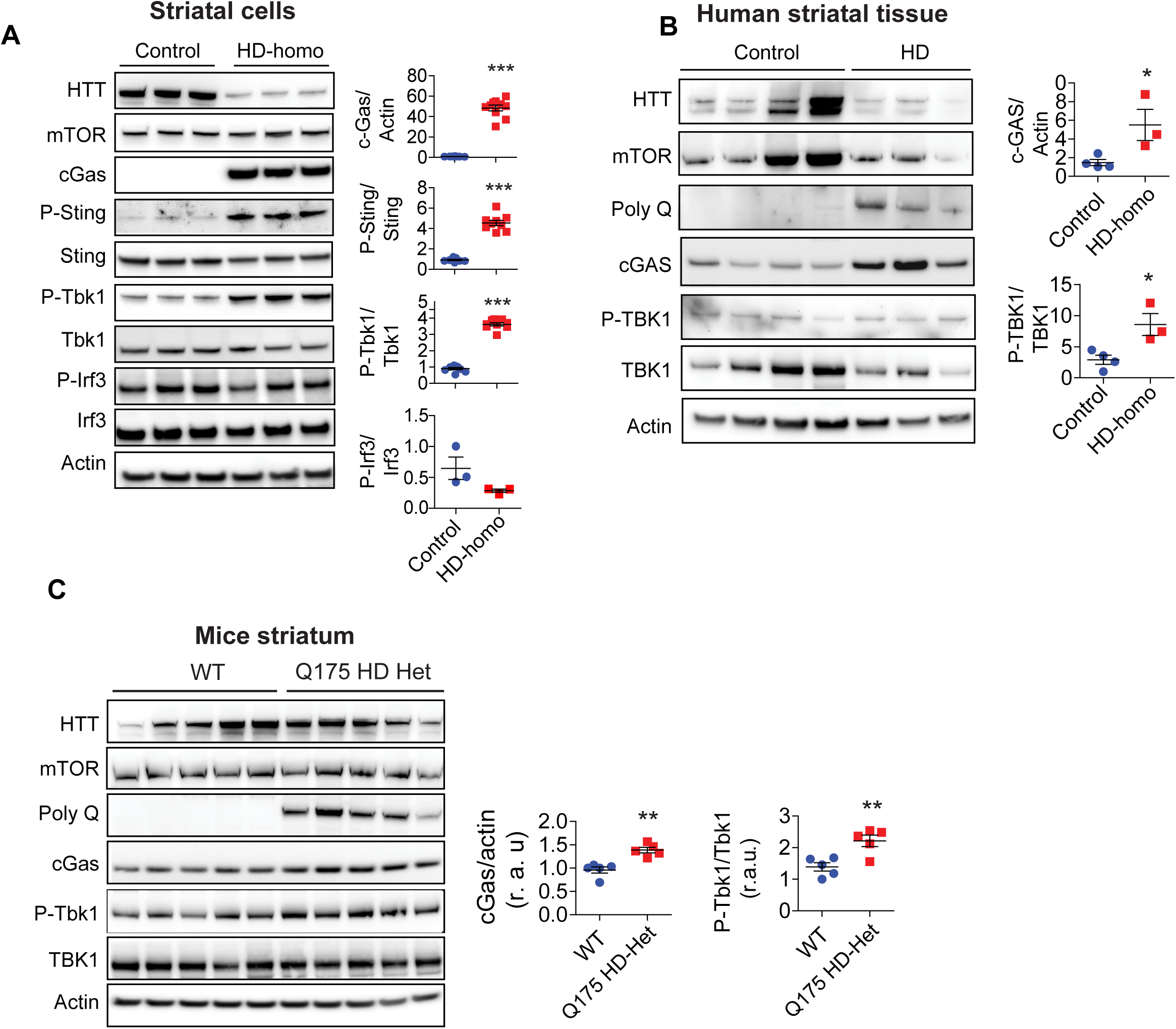
cGAS activity is upregulated in HD, (**A-C**) Representative Western blot for indicated proteins in the striatal neuronal cells (n = 3-8), (A), human unaffected (control, n = 4) and HD affected (n = 3) striatum (B), WT and Q175 HD het (neo-) striatum (n = 5 mice/ group) (C). Error bar represents mean ± SEM, \**p*< 0.05, \*\**p*< 0.01, \*\*\**p*< 0.001 by Student’s-*t* test.

**Figure 4.**
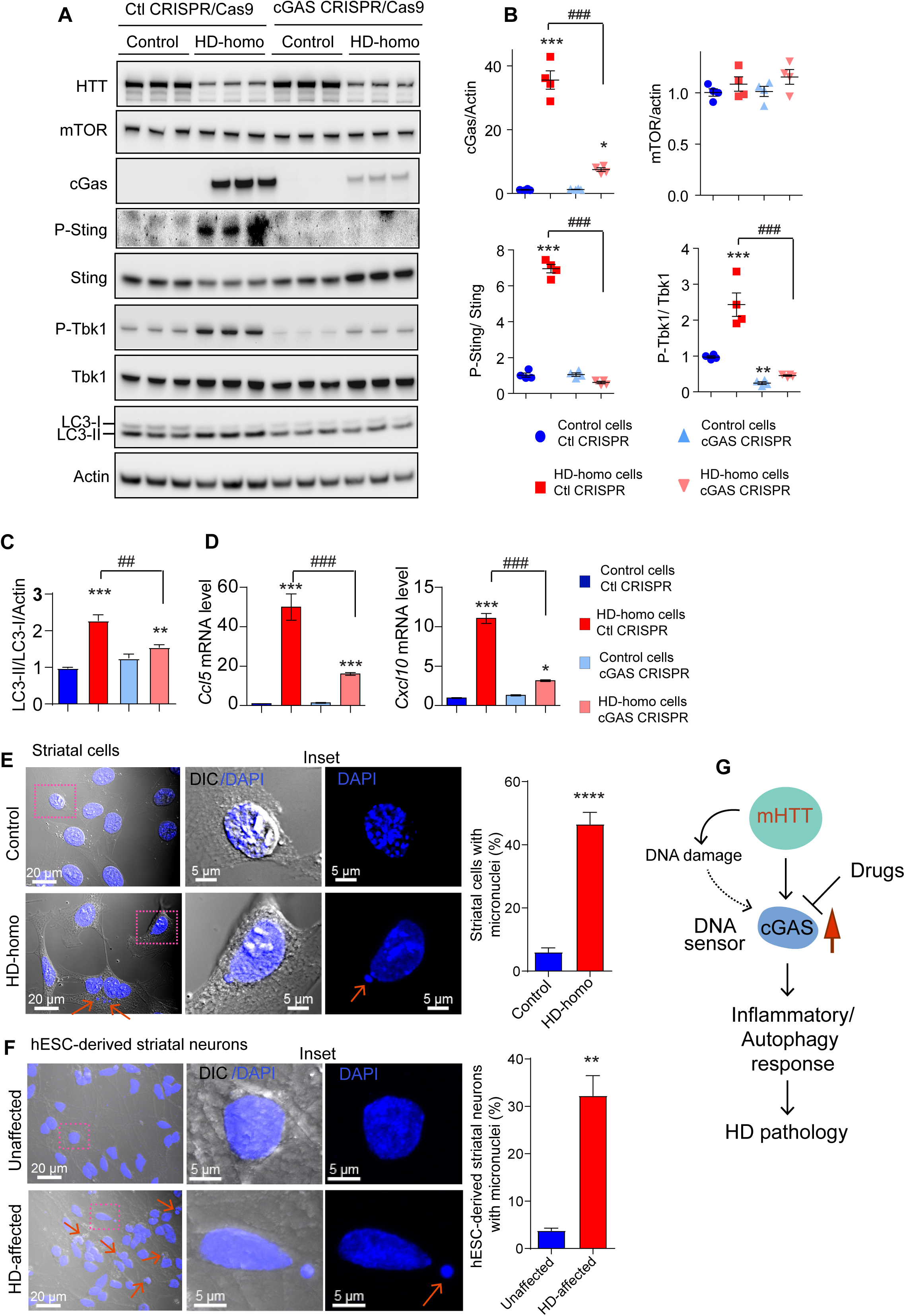
Depletion of cGAS inhibits the upregulation of autophagy and inflammatory response in HD cells. (**A**) Representative Western blot of indicated proteins from control and HD-homo striatal neuronal cells that are generated using control CRISPR/CAS-9 (control) or cGAS gRNA expressing CRISPR/CAS-9 cells (cGAS depleted cells). (**B**) Error bar represents mean ± SEM, (n =4, independent experiments), \**p*< 0.05, \*\**p*< 0.01, \*\*\**p*< 0.001 compared to control cells Ctl CRISPR and ^###^ *p*< 0.001 between HD-homo cells control CRISPR and HD-homo cells cGAS CRISPR by One-way ANOVA test followed by Tukeys post hoc test. (**C**) Indicates the LC-3 protein levels in control and cGAS depleted cells. Error bar represents mean ± SEM, (n = 3, independent experiments), \*\**p*< 0.01, \*\*\**p*< 0.001 compared to control cells Ctl CRISPR and ^##^ *p*< 0.01, between HD-homo cells control CRISPR and HD-homo cells cGAS CRISPR by One-way ANOVA test followed by Tukeys post hoc test. (**D**) qPCR analysis of Ccl5 and Cxcl10 mRNA. Error bar represents mean ± SEM, (n = 3, independent experiments), \**p*< 0.05, \*\*\**p*< 0.001 compared to control cells Ctl CRISPR and ^###^*p*< 0.001, between HD-homo cells control CRISPR and HD-homo cells cGAS CRISPR by One-way ANOVA test followed by Tukeys post hoc test. **(E, F)** Indicates confocal and differential interference contrast (DIC) images of striatal cells (E) and hESC-derived striatal neuron (F). Arrow indicates micronucleus in the cytoplasm. Error bar represents mean ± SEM. \*\**p*< 0.01, \*\*\*\**p*< 0.0001 by Student’s-t-test (control = 308 cells, HD-homo = 134 cells; unaffected = 569 neurons, HD affected = 403 neurons). (**G**) Model depicts mHTT expressing cells increases DNA sensor cGAS levels that controls inflammatory and autophagy responses and thus HD pathogenesis.

### HD cells harbor cytoplasmic micronuclei

We also considered the question of how cGAS is upregulated. A previous study showed that cGAS can be upregulated in the presence of cytoplasmic micronuclei (42), which are discrete DNA fragments that are independent of the main nucleus, a condition which may reflect DNA damage (48). Since we found a rapid upregulation of cGAS levels and its activity in HD cells, we wondered whether there was any evidence of micronuclei in HD cells. To our surprise, we found numerous micronuclei in the HD-homo cells (43% of cells) when compared to the control cells (5%) (**Fig. 4E**). Similarly, HD striatal neuron derived from human embryonic stem cells (hESC) also showed numerous cytoplasmic micronuclei compared to neuron derived from unaffected hESC (Fig. 4F). This data suggests that the presence of micronuclei might be a cause for the upregulation of cGAS and its activity in HD.

Together, our model predicts that mHTT increases cGAS expression, potentially due to cytoplasmic micronuclei, which may in turn affect inflammatory and autophagy responses in HD (**Fig. 4G**).

## Discussion

We found that cGAS levels and activity are upregulated in HD. We discovered using ribosome profiling and biochemical data that cGAS levels are induced and show high ribosome occupancy in a mHTT-copy-number dependent manner (**Fig. 1A and 2A**), indicating that the cGAS expression is CAG dependent. What are the mechanisms by which cGAS is induced in HD and what causes its high ribosome occupancy? Recently, we demonstrated that mHTT stalls ribosomes and inhibits translation elongation [(38), revised], and at the same time increases the protein expression of selected mRNA such as *FMR1*. Because cGAS shows a high ribosome occupancy in exon 1, we speculate that mHTT can directly regulate the translation of *cGAS* mRNA. Alternatively, cGAS may be induced in response to DNA damage/repair, which has been well reported in HD (49-52). Consistent with this, we found numerous micronuclei in the HD cells indicating that cGAS may be induced due to the presence of abnormal nuclei in the cytoplasm (**Fig. 4E, F**). Therefore, one or more mechanisms could be involved in cGAS upregulation in HD, a possibility that needs to be investigated further. Our cGAS depletion study clearly demonstrates that cGAS induction activates the canonical STING/TBK1 pathway and regulates *Cxcl10* and *Ccl5*, but not *IRF3*, expressions in HD striatal cells. Since IRF3 and the related IRF7 are expressed in neuronal cells (53,54), we cannot rule out the possibility that cGAS may induce these targets depending upon the cell types in HD. Moreover, TBK1 has multiple targets that have been previously implicated in HD. For example, TBK1 regulates p62, an autophagy regulator, as well as nuclear factor-kB signaling and mTORC1 signaling, which are also regulated by mHTT (29,30,55-61). This indicates that cGAS signaling may operate at multiple levels in HD. Nevertheless, as mHTT mediates a complex array of important pathways in the cells, such as nutrient-sensing signaling, autophagy, DNA damage, and inflammatory responses, understanding the molecular details of how mHTT intersects with cGAS signaling is critical to further elucidating and eventually interfering with disease-promoting mechanisms. How does cGAS activation might regulate striatal vulnerability in HD? In the striatum, Rhes which increase mHTT solubility and cell-to-cell transport via tunneling-nanotube (TNT)-like membrane protrusions, may further propagate intercellular inflammatory responses (62-66). Collectively, we demonstrate that the cGAS protein and its activity are upregulated in HD, and that depletion of cGAS or its pharmacological inhibition may slow or even halt HD pathogenesis.

## Acknowledgments

We would like to thank; Dr. Long Yan of Max Plank Institute of Neuroscience, Jupiter, Florida, for imaging help. We would like to thank Melissa Benilous for her administrative help and the members of the lab for their continuous support and collaborative atmosphere. This research was partially supported by a training grant in Alzheimer’s drug discovery from the Lottie French Lewis Fund of the Community Foundation for Palm Beach and Martin Counties. This research was supported by funding from National Institutes of Health/National Institute of Neurological Disorders and Stroke grant R01-NS087019-01A1, National Institutes of Health/National Institute of Neurological Disorders and Stroke grant R01-NS094577-01A1, and grants from Cure Huntington Disease Initiative (CHDI) foundation.

## Author contributions

S.S identified cGAS as one of the top high ribosome occupancy targets in ribosome profiling data in HD cells. S.S further conceptualized the project with M.S., who carried out cell culture, biochemical work and analyzed micronuclei. S.R prepared the human embryonic stem cell derived striatal neurons and DAPI staining. S.S prepared all the RFP/mRNA profiles via the UCSC (University of California Santa Cruz) browser. N.S performed Western blotting from HD patient tissue and HD mouse models. N.M.J prepared tissue from HD mouse models and ran Western blot. S.S wrote the paper with input from co-authors.

## Materials and methods

### Cell culture

Mouse STHdh^Q7/Q7^ (control), STHdh^Q7/Q111^ (HD-het), and STHdh^Q111/Q111^ (HD-homo) striatal neuronal cells (37). Striatal cells were cultured in growth medium containing Dulbecco’s modified Eagle’s medium high glucose (Thermo Fisher Scientific) with 10% fetal bovine serum (FBS), 1% penicillin-streptomycin at 33°C and 5% CO_2_, as described in our previous works (61,64,67).

### hESC-derived neuron culture

Unaffected (Genea-019) and HD-affected (HTT-48Q, Genea-020) hESC lines were obtained from CHDI. The lines were grown and expanded in feeder-free conditions on vitronectin using Essential 8 plus medium (Thermo Fisher Scientific) at 37°C and 5% CO_2_. Medium was replaced daily. hESCs were differentiated into striatal neurons using a previously described protocol (68) Briefly, on Day 0, hESCs were detached with Versene to form embryoid bodies (EBs) in Neural induction medium (NIM, DMEM/F12: Neurobasal plus (1:1), 1x N2 supplement, 1x B27 supplement, 1x GlutaMAX, 1x penicillin streptomycin) plus LDN-193189 (100 nM, Sigma), SB431542 (10 µM, Sigma), XAV939 (4 µM, Selleck), SAG (100 nM, Selleck). The EBs were then plated in neural proliferation medium (NPM, DMEM/F12: Neurobasal plus (1:1), 0.5x N2 supplement, 1x B27 supplement, 1x GlutaMAX, 1x penicillin streptomycin) plus LDN-193189 (100 nM), SB431542 (10 µM), XAV939 (4 µM), SAG (100 nM) onto plastic plates coated with polyornithine (15 µg/ml, Sigma)/laminin (5 µg/ml, Sigma)/fibronectin (5 µg/ml, Thermo fisher Scientific) on Day 4. Next day the medium was changed to NPM medium plus LDN-193189 (100 nM), SB431542 (10 µM). Medium was changed every other day. On Day 9, medium was changed to NPM medium without any small molecules. Within days 11–16 of differentiation, the cells were dissociated with Accutase and replated onto polyornithine/fibronectin/laminin-coated coverslips in neural differentiation medium (NDM, Neurobasal plus, 1x B27 supplement, 1x GlutaMAX, 1x penicillin streptomycin) plus ascorbic acid (AA, 0.2 mM, Sigma), and DAPT (1µM, Tocris), BDNF (20 ng/ml, Prospec), GDNF(10 ng/ml, Prospec). The medium was changed every other day with NDM medium supplemented with AA (0.2 mM), and DAPT (1µM), BDNF (20 ng/ml), GDNF (10 ng/ml). The differentiated striatal neurons on Day 22 were used for staining with DAPI.

### Ribosome Profiling

RNAase foot printing, generation of cDNA libraries from ribosome protected mRNAs, generation of cDNA libraries and sequencing, generation of mRNA-seq libraries, Ribo-Seq, RNA-seq quality control and mapping the reads to UCSC browser and ribosome pause analysis, were carried out as described in our previous work (38).

### Mice

C57BL/6J (WT) and the Q175DN HD mouse (B6J.zQ175DN KI #029928) obtained from Jackson laboratories and maintained in our animal facility according to Institutional Animal Care and Use Committee at The Scripps Research Institute. Three to four months old mice were used for the experiment.

### Antibodies

Huntingtin (MAB2166) antibody were obtained from, Millipore-Sigma. Actin (sc-47778) and cGAS (sc-515777) were purchased from Santacruz Biotechnology. Huntingtin (5656), mTOR (2983), STING (50494), phospho-STING Ser365 (72971), cGAS (31659), TBK1 (3504) and phospho-TBK1 Ser172 (5483), IRF3 (4302) and phosphor-IRF3 Ser396 (29047) antibodies were purchased from Cell Signaling Technology. HRP-conjugated secondary antibodies were from Jackson ImmunoResearch Inc.

### Generation of cGAS depleted striatal cells

cGAS depleted striatal cells were generated using cGAS CRISPR/Cas9 plasmids from Santa cruz Biotechnologies. First, we transfected the striatal neuronal cells (control and HD-homo) with cGAS CRISPR/Cas9 plasmid ((sc-437363)) or CRISPR/Cas9 control plasmid (SC-418922) in 10cm dish. After 48 h we sorted the cells based on GFP fluorescence and re-cultured them. We passaged them 2-3 time and prepared lysate to confirm the cGAS depletion (∼90%) by Western blotting using cGAS antibody.

### Protein expression and Western blots

To check the protein expression level striatal neuronal cells (control and HD) were plated in 6 well plate (2×10^5^ cells per well). After 24 hours, cells were washed in PBS and lysed in buffer containing 1% Triton X-100 in 50 mM Tris-HCl, pH 7.5, 150 mM NaCl and 1x protease inhibitor cocktail (Roche, Sigma) and 1x phosphatase inhibitor (PhosStop, Roche, Sigma), sonicated for 3 × 5 sec at 20% amplitude, and cleared by centrifugation for 10 min at 11,000g at 4°C. Protein concentration was determined with a bicinchoninic acid (BCA) protein assay reagent (Pierce). Equal amounts of protein (30-40 µg) were loaded and were separated by electrophoresis in 4 to 12% Bis-Tris Gel (Thermo Fisher Scientific), transferred to polyvinylidene difluoride membranes, and probed with the indicated antibodies. HRP-conjugated secondary antibodies (Jackson ImmunoResearch Inc.) were probed to detect bound primary IgG with a chemiluminescence imager (Alpha Innotech) using enhanced chemiluminescence from WesternBright Quantum (Advansta)(67). The band intensities were quantified with ImageJ software (NIH). Phosphorylated proteins were then normalized against the total protein levels (normalized to actin). Mice striatal tissues were homogenized in radioimmunoprecipitation assay (RIPA) buffer [50 mM Tris-HCl (pH 7.4), 150 mM NaCl, 1.0% Triton X-100, 0.5% sodium deoxycholate, 0.1% SDS,] with 1x complete protease inhibitor cocktail (Roche, Sigma) and 1x phosphatase inhibitor (PhosStop, Roche, Sigma), followed by a brief sonication for 2 × 5 sec at 20% amplitude and cleared by centrifugation for 10 min at 11,000g at 4°C. Protein estimation was done using a BCA method and proceeded to Western Blotting as mentioned above.

### Human HD patient samples

Human brain tissue (Caudate nucleus) samples of grade 1 HD-affected patient (HSB # 3358, 2706, 3744), grade 2 HD-affected patient (HSB # 2858, 3432, 3635, 3872, 4072, 3159), grade 3 HD-affected patient (HSB # 4344, 2869, 2972, 4518, 4254), grade 4 HD-affected patient (HSB # 5078, 2903) and normal donor controls (HSB # 4615, 4823, 5293, 4340, 4135) were obtained from the Human Brain and Spinal Fluid Resource Center VA West Los Angeles Healthcare Center, Los Angeles, CA. Human tissue was homogenized in lysis buffer [50 mM tris (pH 7.4), 150 mM NaCl, 10% glycerol, and 1.0% Triton X-100] with protease and phosphatase inhibitors, followed by a brief sonication for 6 s at 20% amplitude and cleared by centrifugation for 10 min at 11,000g at 4°C. Protein estimation was done using a BCA method and proceeded to Western Blotting as mentioned above.

### cDNA preparation and Real-time PCR

RNA was extracted from the fractionated samples following lysis in Trizol reagent. 250 ng RNA was used to prepare cDNA using Takara primescripttm kit (Cat no. 6110A) using random hexamers. The qRT-PCR of genes was performed with SYBR green (Takara RR420A) reagents. Primers for all the genes were designed based on sequences available from the Harvard qPCR primer bank. The primer sequences are as follows:

*Gapdh* mouse (Forward primer) 5’ primer AGGTCGGTGTGAACGGATTTG (Reverse primer) 3’ primer TGTAGACCATGTAGTTGAGGTCA

*Ccl5* mouse (Forward primer) 5’ primer GCTGCTTTGCCTACCTCTCC (Reverse primer) 3’ primer TCGAGTGACAAACACGACTGC

*Cxcl10* mouse (Forward primer) 5’ primer CCAAGTGCTGCCGTCATTTTC (Reverse primer) 3’ primer GGCTCGCAGGGATGATTTCAA

*Gapdh* human (Forward primer) 5’ primer GGAGCGAGATCCCTCCAAAAT (Reverse primer) 3’ primer GGCTGTTGTCATACTTCTCATGG

*Ccl5* human (Forward primer) 5’ primer CCAGCAGTCGTCTTTGTCAC (Reverse primer) 3’ primer CTCTGGGTTGGCACACACTT

### Staining with DAPI

Striatal neuronal cells (Control and HD) plated in glass bottom dishes or hESC-derived striatal neurons on coverslips were washed in D-PBS and fixed for 10 min in 4% PFA (Electron Microscopy Sciences). The cells were washed 3 times with D-PBS and incubated with DAPI (4′,6-diamidino-2-phenylindole) for 10 min.

### Image processing and micronuclei quantification

All the fluorescent confocal images were taken in Zeiss 880 microscope using 63X oil immersion Plan-apochromat objective (1.4 NA). Excitation was via a 405 nm diode-pumped solid-state laser. Pinholes were set so that the section thickness was equal for all channels and ≤ 1?AU. Images were acquired with an optimal Z-step of 0.27 µm covering the whole cellular volume. Processing was performed with Zen software black/blue edition 2012. For micronuclei quantification, we counted the cells with small DAPI positive puncta separated from nucleus.

### Statistical analysis

Unless otherwise noted all experiments were carried out in duplicates and repeated at least three times. The statistical comparison was carried out between groups using one-way ANOVA followed by Tukey’s multiple comparison test or Student’s *t*-test, and significance values were set at p < 0.05, using Graphpad Prism 7.

